# Separation and identification of permethylated glycan isomers by reversed phase nanoLC-NSI-MS^n^

**DOI:** 10.1101/2020.08.04.236349

**Authors:** Simone Kurz, M. Osman Sheikh, Shan Lu, Lance Wells, Michael Tiemeyer

## Abstract

High performance liquid chromatography has been employed for decades to enhance detection sensitivity and quantification of complex analytes within biological mixtures. Among these analytes, glycans released from glycoproteins and glycolipids have been characterized as underivatized or fluorescently tagged derivatives by HPLC coupled to various detection methods. These approaches have proven extremely useful for profiling the structural diversity of glycoprotein and glycolipid glycosylation but require the availability of glycan standards and secondary orthogonal degradation strategies to validate structural assignments. A robust method for HPLC separation of glycans as their permethylated derivatives, coupled with in-line MSn fragmentation to assign structural features independent of standards, would significantly enhance the depth of knowledge obtainable from biological samples. Here, we report an optimized workflow for LC-MS analysis of permethylated glycans that includes sample preparation, mobile phase optimization, and MS^n^ method development to resolve structural isomers on-the-fly. We report baseline separation and MS^n^ fragmentation of isomeric N- and O-glycan structures, aided by supplementing mobile phases with Li^+^, which simplifies adduct heterogeneity and facilitates cross-ring fragmentation to obtain valuable monosaccharide linkage information. Our workflow has been adapted from standard proteomics-based workflows and, therefore, provides opportunities for laboratories with expertise in proteomics to acquire glycomic data with minimal deviation from existing buffer systems, chromatography media, and instrument configurations. Furthermore, our workflow does not require a mass spectrometer with high-resolution/accurate mass capabilities. The rapidly evolving appreciation of the biological significance of glycans for human health and disease requires the implementation of high-throughput methods to identify and quantify glycans harvested from sample sets of sufficient size to achieve appropriately powered statistical significance. The LC-MSn approach we report generates glycan isomeric separations, robust structural characterization, and is amenable to auto-sampling with associated throughput enhancements.

## INTRODUCTION

Every living cell is covered in a dense and complex array of glycoconjugates that modulate various biological processes such as molecular recognition, cell-cell, and cell-matrix interactions. The structural diversity of glycans arises from differences in monosaccharide compositions, anomeric confirmation, glycosidic linkages, branching, and their attachment to protein or lipid (1). A hallmark of protein glycosylation is the phenomenon of microheterogeneity, or the property that any single glycosylation site may be unmodified or modified by more than one glycan structure when comparing molecules of the same protein, even if produced in the same cell type. This diversity presents unique challenges for characterizing glycoprotein glycosylation and creates vital needs for technologies that robustly capture glycan structural details (2). The biochemical and analytical tools currently deployed for glycomic analysis are almost as diverse as the biological roles of glycans and are highly influenced by the research purpose.

Given the complexity of glycan structures, with special reference to the broad range of isomeric possibilities, glycan separation by liquid chromatography (LC) has emerged as a state-of-the-art approach in glycomics. Almost all previous LC work has been done on underivatized or fluorescently-tagged glycans, allowing high sensitivity of detection and high resolution separations based on size, charge, and hydrophobicity (3). These approaches require previously characterized glycan standards as reference materials or sequential enzymatic (e.g. exoglycosidase sequencing) and/or chemical treatments (e.g. hydrofluoric acid) for complete glycan characterization. While offline or online mass spectrometry (MS) based methods can be combined with tagged-glycan separations, the sensitivity and information content ofthe resulting data is significantly influenced by the ionization characteristics and chemical nature ofthe tag. In that respect, permethylation has emerged as a routine derivatization procedure that improves sensitivity by enhancing MS ionization. Glycan permethylation converts the oxygen of all hydroxyl groups, including the sialic acid carboxylates, into an O-methyl groups (O-CH_3_). Because glycans generally have a large number of free hydroxyls, the O-methyl groups of permethylated glycans dominate the chemical behavior of the molecule, enhancing and equalizing ionization across diverse glycan structures (4). Permethylation thereby allows for the analysis and quantification of acidic and neutral structures in the mass spectrometer’s positive ion detection mode.

In sequential fragmentation MS/MS and MS^n^ analysis, ‘scars’ of a free hydroxyl group instead of O-methyl are detected at the original position of glycosidic linkages, thereby providing valuable linkage information through cross-ring cleavages. Permethylation and heavy metal cations facilitate these cross-ring fragments enabling in-depth structural characterization of glycans by direct infusion into the mass spectrometer without LC separation (5-8). However, structural elucidation of isomeric permethylated glycans poses analytic challenges in direct infusion MS since they are detected at the same m/z. This complication can be conquered by combining LC separation of permethylated glycans with intelligent MS^n^ in realtime. Various LC-MS methods for analyzing permethylated glycans have been published recently (9, 10). Here, we report our solution, which is designed to be rapidly adaptable by laboratories familiar with standard proteomic workflows. Using an Ultimate 3000 RSLC HPLC system coupled to a Velos Pro Dual Pressure Linear Ion Trap we describe parameters of sample preparation, mobile phase and gradient optimization, as well as post-acquisition data analysis of permethylated chemically synthesized standards and glycans released from either standard proteins or biological sample mixtures. We have achieved baseline separation and MS^n^ fragmentation of isomeric N- and O-glycan structures to aid in confident identification and quantification by pushing the ionization into a single adduct form, namely lithium. Our workflow simplifies post-acquisition data analyses and enhances throughput of glycomic data collection without drastically changing chromatography media used for proteomics or requiring a mass spectrometer with high-resolution/accurate mass capabilities.

## EXPERIMENTAL PROCEDURES

### Materials and Reagents

PNGase F (N-glycanase) was obtained from the Complex Carbohydrate Research Center, University of Georgia, (Dr. Kelley Moremen). PNGase A was acquired from NEB. N-Glycan standards were obtained from the Consortium for Functional Glycomics (CFG) and from The Scripps Research Institute (Dr. James Paulson). O-Glycan standards were obtained as part of an NIH Common Fund grant (R21AI123161) from the University of Georgia (Dr. Christopher M. West). Sodium hydroxide (50%) was purchased from Fisher Scientific. Sep-Pak C18 disposable extraction columns were obtained from Waters Corporation (Milford, MA, USA). AG-50W-X8 cation exchange resin (H^+^ form) was purchased from Bio-Rad and trifluoroacetic acid from Pierce. Malto-series oligosaccharides were obtained from Wako Chemicals (Richmond, VA). Trypsin, Chymotrypsin, bovine pancreatic ribonuclease B (RNaseB), Fetuin, Dextran (Leuconostoc spp., Cat# 31388) and all other chemical reagents were purchased from Sigma-Aldrich (St. Louis, MO, USA). Fly powder, a dried preparation of *Drosophila* embryo glycoproteins, was prepared as previously described and used as a source of pauci-mannose glycans (11). Mouse brain extracts were prepared as previously described for analysis of brain O-glycans (12). HIV gp120 was recombinantly expressed in glyco-engineered *Pichia pastoris* strains obtained from GlycoFi (Lebanon, NH) in order to generate proteins with restricted glycan diversity (13, 14).

### Preparation ofprotein rich powderandsubseguent O-glycan release

Primary mouse tracheal epithelial cells (mTEC), mouse brain (Pel-Freez Biologicals), or HEK293 cells (kind gift from Dr. Henrik Clausen, University of Copenhagen, Denmark) were homogenized in ice-cold 50% (v/v) aqueous methanol and delipidated with chloroform/methanol/water (4:8:3, v/v/v) as described previously (11, 15). Insoluble proteins were precipitated by centrifugation and protein pellets were washed twice with ice-cold acetone prior to drying under a gentle Nitrogen stream to produce fine protein powder. 2-3 mg of protein powder were then subjected to reductive β-elimination and the released O-glycan alditols purified as described previously (16). Briefly, protein powder was resuspended in 100 mM NaOH containing 1 M NaBH_4_ and incubated for 18 h at 45°C in a glass tube sealed with a Teflon-lined screw top. After incubation, the reaction mixture was neutralized with 10% acetic acid on ice and desalted using a AG-50W-X8 (H^+^ form) column (1 mL bed volume) prior to borate removal and Sep-pack C18 cartridge clean-up.

### Release and reduction of N-qlycans

Preparation of glycopeptides and release of N-glycans was performed as described previously (11). Briefly, 20-75 *μ*g of glycoproteins (RNAse B, Fetuin, gp120) were dried and resuspended in trypsin buffer (0.1 M Tris-HCl, pH 8.2, containing 1 mM CaCl_2_) by sonication and boiled for 5 min before addition of trypsin and chymotrypsin solutions. After incubation for 18 h at 37 °C, the (glyco)peptide mixture was boiled for 5 min and adjusted to 5% AcOH (acetic acid) prior to a Sep-Pak C18 cartridge column clean up. Glycopeptides were eluted stepwise in 20% isopropyl alcohol in 5% acetic acid and 40% isopropyl alcohol in 5% acetic acid. Both, the 20 and 40% isopropyl alcohol fractions were pooled and evaporated to dryness. Dried glycopeptides were resuspended in 25 mM sodium phosphate buffer, pH 7.5, for digestion with PNGaseF (fetuin, ribonuclease B) or in 50 mM ammonium acetate buffer, pH 4.5, for digestion with PNGaseA (fly powder) prior to incubation for 18 h at 37 °C. PNGase-released oligosaccharides were separated from residual (glyco)peptides by another round of Sep-Pak C18 cartridge clean-up.

For the reduction of PNGase-released N-glycans and standards, dried glycans were resuspended in 100 *μ*L of 50 mM NaOH, vortexed and ultrasonic water bath sonicated prior to the addition of 100 *μ*L 2% (w/v) NaBH_4_ in 50 mM NaOH. After incubation for at least 4 hours at RT, 200 *μ*L of 10% (v/v) AcOH were added to neutralize the reaction. Samples were dried down under a gentle Nitrogen stream prior to borate removal by adding 400 *μ*L of 10% (v/v) AcOH in methanol and repeated evaporation (up to 3 times).

### Permethylation

All released N- and O-linked glycans as well as standards were permethylated prior to MS analysis according to the method by Anumula and Taylor (17). Dextran, malto-series oligosaccharide standards and CFG N-glycan standards were permethylated with ^13^C methyliodide (^13^C-Mel).

### nanoLC-MS/MS ofPermethylated Glycans

Dried permethylated glycans and standards were dissolved in 100% methanol, and an aliquot (usually 5 to 10% of the total sample) was combined with an internal standard mix (^13^C-permethylated DP4) and mobile phase A. For each nanoLC-MS/MS analysis, 3-6 μL of the prepared sample mix was injected for liquid chromatography separation at 60°C and a constant flow rate of 300 nL/min using an Ultimate 3000 RSLC (Thermo Fisher Scientific/Dionex) equipped with a PepMap Acclaim analytical C18 column (75 μm x 15 cm, 2 μm pore size) coupled to a Thermo Fisher Scientific Velos Pro Dual-Pressure Linear Ion Trap mass spectrometer, ending with sample ionization via a stainless steel emitter. The three solvent systems used were either 0.1% formic acid in water for mobile phase A and 0.1% formic acid in 80% acetonitrile for mobile phase B (“Proteomics buffers”); 0.1% acetic acid containing 0.1 mM LiOAc for mobile phase A and 0.1% AcOH in 80% ACN containing 0.1 mM LiOAc for mobile phase B (“Low lithium buffers”); or 0.02% acetic acid containing 1 mM LiOAc for mobile phase A and 0.02% AcOH in 80% ACN containing 1 mM LiOAc for mobile phase B (“High lithium buffers”).

For O-glycans, after equilibrating the column in 99% mobile phase A for 5 mins, separation was achieved using a linear gradient from 30% to 70% mobile phase B over 150 mins. For N-glycans, after equilibrating the column in 99% mobile phase A for 5 mins, separation was achieved using a linear gradient from 45% to 70% mobile phase B over 150 mins. The analytical column was regenerated after each run by ramping to 99% mobile phase Bfor 10 minutes and then returning to 99% mobile phase A to re-equilibrate.

The Velos Pro Dual Pressure Linear Ion Trap Mass Spectrometer was operated in positive ion mode with a spray voltage of 1.8-2.2 kV and capillary temperature set at 210°C. The MS method consisted of first collecting a Full ITMS (MS1) survey scan in profile mode and m/z mass range from 500-2000 with automatic gain control (AGC) Target set to 30000.00, followed by data-dependent MS2 fragmentation of the Top 3-5 most intense peaks using CID at 40-42% collision energy and an isolation window of 2 m/z. Dynamic exclusion parameters were set to exclude ions for fragmentation for 15 seconds if they were detected and fragmented 5 times in 15 secs. For targeted MS^n^ approaches, intelligent fragmentation methods were generated using neutral loss function and product dependent MS^3-5^ acquisition to determine sialic acid linkage of terminally sialylated N-glycan structures. To normalize elution times on the RP-column in terms of glucose units (g.u.), ^13^C-permethylated dextran (4-10 g.u.) was analyzed before and after a sequence of samples. After the completion of a sample set, in preparation for limited HPLC use for an extended period of time, the post-column fused silica lines were manually flushed with 50% methanol in water to avoid clogging of the stainless steel emitter with lithium salts. All data were processed manually using the Xcalibur software package 2.0. GRITS Toolbox, a freely available tool for semi-automated annotation of glycomic MS data was also employed to generate candidates for structural assignment (18).

## RESULTS

### Sample Preparation

Our workflow can be applied to any biological starting material. In the first preparation steps, glycosphingolipids are extracted from the biological sample prior to acetone precipitation to obtain protein rich powder (**Figure 1**). Starting with 1-2 mg of protein powder each, we can prepare N- and O-glycans. For N-glycans, the glycoproteins are proteolytically cleaved into glycopeptides (usually by trypsin and/or chymotrypsin) and purified using a C18 solid-phase extraction column. N-glycans are then enzymatically released with PNGase F. For LC-NSI-MS analysis, released N-glycans can also be reduced in mild sodium borate conditions prior to borate removal and permethylation for improved chromatographic separation (see **Supplemental Figure 1**). O-glycans are chemically released by reductive β-elimination, followed by desalting, borate removal and C18 clean up. Following permethylation and phase partitioning, we subjected the organic phase, which contains neutral and non-sulfated glycans, to analysis by LC-NSI-MS using various buffer and gradient conditions.

**Figure 1.**
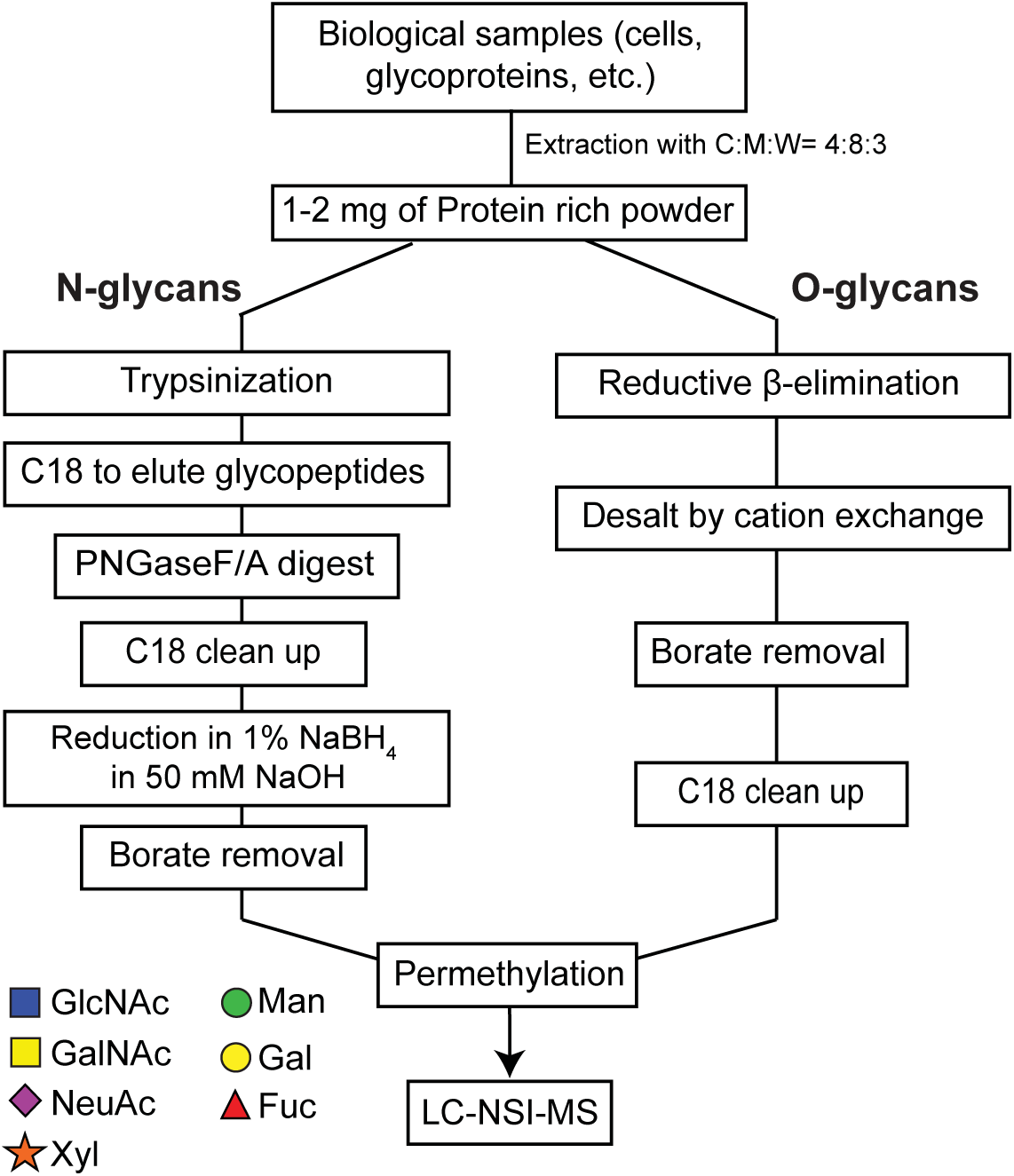
General workflow for preparation of N- and O-glycans. Using virtually any biological source as starting material, protein-rich powder can be generated via organic delipidation as described previously (11). N-linked glycans can be released enzymatically (PNGase F or PNGase A) followed by mild reduction of the free reducing terminus. O-linked glycans can be released chemically via reductive β-elimination. Released glycans of either class are subjected to permethylation followed by direct analysis by LC-NSI-MS. Symbol and Text nomenclature for representation of glycan structures is displayed according to the Symbol Nomenclature for Glycans (SNFG) (39).

### O-Glycan Separation and Mobile Phase Optimization

In order to test the separation of permethylated O-glycans using our platform, initial RP C18 nanoLC-separations were performed on O-glycans released by reductive beta-elimination from primary mouse tracheal epithelial cells (mTEC). These initial separations used conventional acidic proteomic mobile phases at an elevated column oven temperature (60 °C) to facilitate baseline separation (**Figure 2, A**). As has been previously described and also in our analysis, lower column oven temperatures (40 °C and 50 °C) resulted in significant peak broadening (10). By generating extracted ion chromatograms (EICs) of expected glycan species, the major mTEC O-glycans were resolved and multiple EIC peaks were identified for some, revealing the possibility of isomeric structures (**Figure 2, B**).

**Figure 2.**
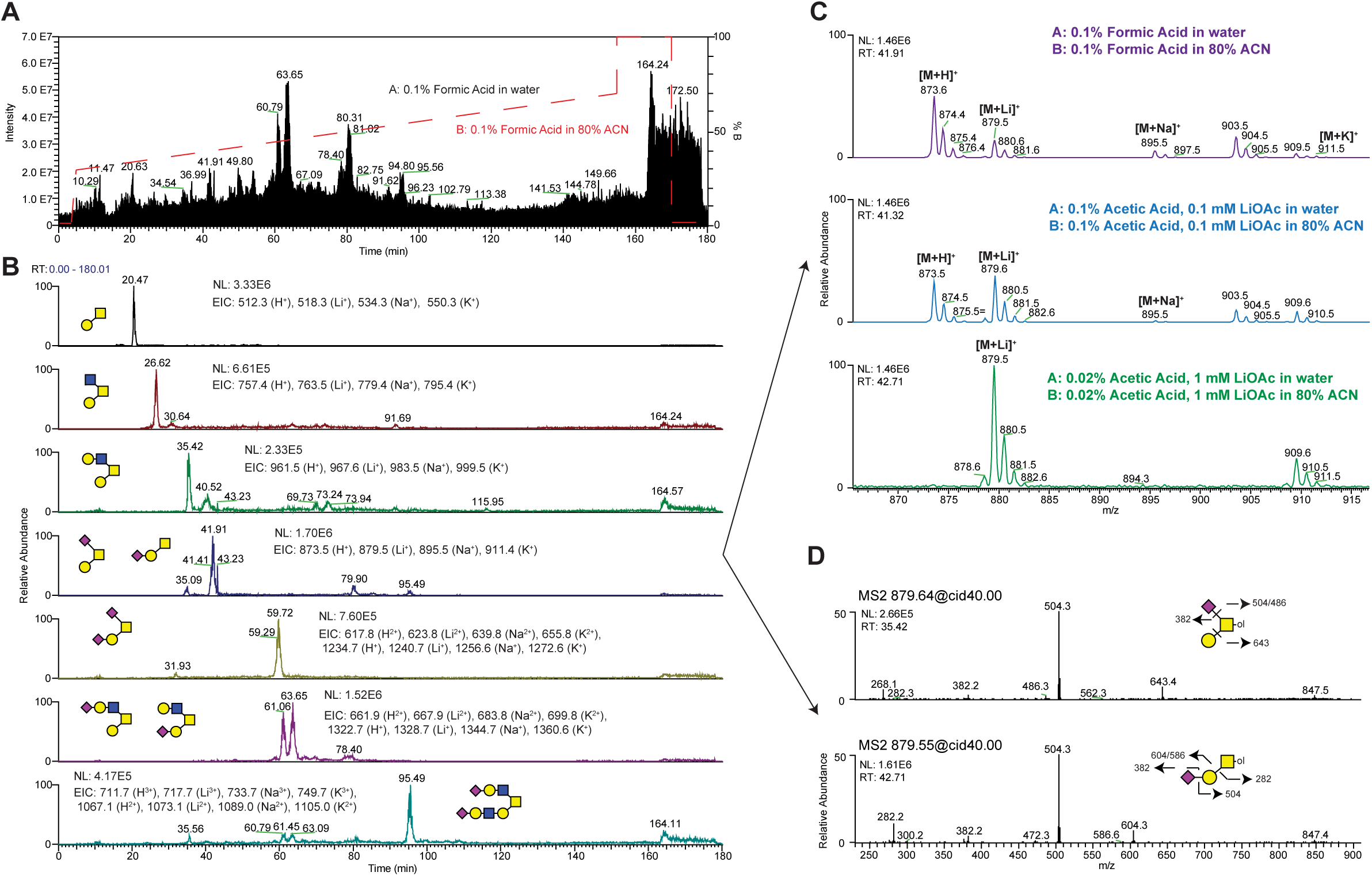
Addition of lithium acetate to LC-mobile phases enhances ionization. Reduced O-glycans from mouse tracheal epithelial cells (mTECs) were separated on a PepMap Acclaim C18 column (75 μm x 15 cm, 2 μm pore size) using three different mobile phase pairs (Buffers A and B) on a Dionex UltiMate 3000 coupled to Thermo Velos Pro ITMS. (**A**) The total current chromatogram and extracted ion chromatograms (**B**) demonstrate baseline separation of permethylated O-glycans using common proteomics buffers (Formic acid/Acetonitrile). (**C**) Observed alkali metal adduct ions for the three mobile phase conditions tested demonstrated optimal ionization and adduct formation in 0.02% Acetic Acid containing 1 mM lithium acetate. (**D**) MS/MS fragmentation oftwo glycan isoforms are shown.

One disadvantage of using acidic mobile phases that are standard for proteomic analyses is the formation of multiple alkali metal glycan adducts which makes data analysis and, importantly, quantification more difficult. As demonstrated for a simple sialyl Tn O-glycan structure, multiple ions including protonated [M+H]^+^, lithiated [M+Li]^+^, sodiated [M+Na]^+^ and potassiated [M+K]^+^ adducts were detected in a Full MS scan using conventional proteomics mobile phases (**Figure 2, C**). In order to promote the ionization into only one molecular ion species and to enhance cross-ring cleavages, lithium acetate was supplemented in the mobile phases at 0.1 mM resulting in an improvement but not complete shift towards lithiated ion species. By increasing the concentration to 1 mM lithium acetate and reducing acid concentration, adduct ions were shifted to essentially lithiated-only forms (**Figure 2, C**). Combining EICs with the data-dependent MS/MS data acquisition in real-time, the structures of two isomeric sialyl Tn O-glycans were resolved with the minor branched structure eluting earlier (35 min) than the more abundant linear structure (42 min) (**Figure 2, D**).

### N-Glycan Separation and Gradient Optimization

To assess the feasibility of separating permethylated N-glycans, we used PNGaseF-released N-glycans from RNAseB as our glycan source. Initial peak-splitting during the LC-separation was observed, presumably due to mutarotation of the anomeric configuration at the free reducing end prior to permethylation (α- vs. β-anomer; **Supplement Figure 1A**). To ameliorate this complication, reduction of PNGase-released N-glycans prior to permethylation is strongly recommended for simplifying glycans to elute as one peak rather than two α- and β-anomer peaks (**Supplement Figure 1B**). Peak splitting was not observed in our mTEC O-glycan analysis since the reducing termini of the released O-glycans are reduced during the release by reductive β-elimination as part of our standard workflow (**Figure 1**).

Using an initial separation gradient of 30% to 70% mobile phase B over 150 mins, baseline separation was achieved for reduced and permethylated paucimannosidic, high-mannose and complex type N-glycan structures released from fly protein powder, RNAseB and Fetuin, respectively (**Figure 3, A**). As this gradient might be insufficient for separation of larger N-glycans such as tetra-antennary structures, a modified gradient with a rapid increase in the first 5 minutes to 45% B followed by a shallow gradient of 45% to 70%B over 150 minutes was applied and was observed to not change the overall elution pattern of the tested N-glycans (**Figure 3, B**). Using our platform and elution conditions, attempts to shorten the gradient to less than 180 minutes as described by others resulted in suboptimal separation of glycans and increased numbers of coeluting species (3, 9, 10).

**Figure 3.**
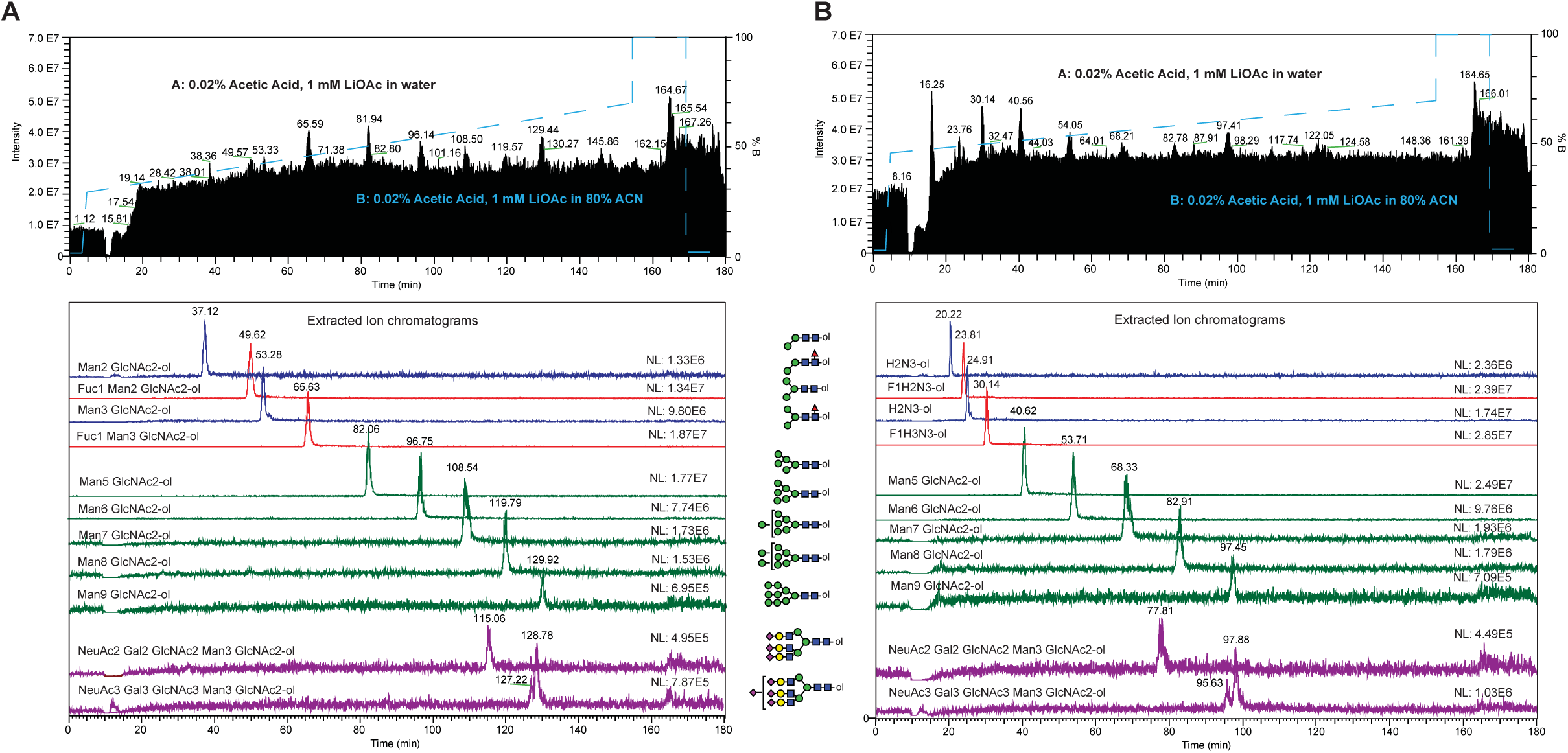
LC-gradient optimization for reduced N-glycans. Initial separation of paucimannose, high-mannose and complex type N-glycans was achieved with a gradient from 30-70% B (**A**), but separation of larger N-glycans (e.g. tetra-antennary) might be insufficient. (**B**) Rapid increase in the gradient to 45% B in 5 min followed by a shallow gradient from 45-70% B seems to be more suitable for larger N-glycans (e.g. tri/tetra-antennary). Attempts to shorten the gradient program to less than 180 min using steeper gradients resulted in suboptimal separation of glycans and co-eluting species.

### Chromatographic retention times can be converted to glucose units

In many instances, the combination of LC retention time and accurate mass of a detected analyte can lead to structural assignment by reference to well-characterized standards. However, assignment/annotation solely based on retention time and mass is frequently impractical due to technical challenges associated with variability in column performance, clogged tubing, clogged emitters, etc.. Therefore, a more robust relationship between glycan structure and retention time can be achieved by referencing glycan elution positions to a hexose polymer ladder, yielding retention times in glucose units or g.u. (19, 20). Dextran polymers ranging in length (degree of polymerization, DP) from 4 to 10 monosaccharide units where reduced and permethylated with ^13^C-Mel to produce a dextran reference ladder of g.u. standards (DP4-10). Aliquots of the reference ladder were injected before and after sample acquisition queues, and the resulting retention times were averaged from EICs for each predicted mass within 10 ppm (**Figure 4, A**). To test the robustness of our method for characterizing the elution positions of glycans with greater complexity than the dextran ladder, we analyzed four CFG standards (107, 108, 109 and 110) which represent bisected, complex branched N-glycans and one CFG standard (121), a sialylated biantennary N-glycan. Separation of all five reduced and ^13^C-permethylated CFG standards was achieved resulting in standard 107 (bisected, biantennary N-glycan) eluting before standard 108 (bisected, triantennary N-glycan with two GIcNAc residues attached to the core a1,3 mannose). Interestingly, a gap of 6 minutes in retention time was detected between the elution of the isomeric structure of standard 108, which is standard 109 (bisected, triantennary N-glycan with two GIcNAc residues attached to the core a1,6 mannose) demonstrating the power of our modified gradient to achieve separations based on size and hydrophobicity (**Figure 4, B**). CFG standard 110 (bisected, tetra-antennary N-glycan) eluted approximately 2.6 minutes after standard 109 and before the final sialylated, biantennary N-glycan standard 121. Using the linear regression fitting from the dextran ladder runs before and after the sample queue, the observed retention times were converted to reverse phase glucose units (RP g.u.) in order to express the elution positions of the analyzed CFG standards (**Figure 4, C**). Dextran ladder was analyzed before a large sample acquisition queue and subsequently after 25 days of continuous operation (**Supplement 2**). Retention times of isomaltose polymers after nearly one month of continuous use were consistent within 2-3%. Furthermore, retention times were consistent regardless of amount injected, in a dynamic range of 15.6 to 1000 fmol of injected standard (**Supplement 3**). Similar to the N-glycan standards, we tested the ability of our method to separate chemically synthesized, reduced and ^13^C-permethylated O-glycan isomeric standards derived from the human pathogen *Trypanosoma cruzi*, the causative agent for Chagas disease in South America (21,22). The four tetrasaccharide O-glycan isomers analyzed only differ by either the cyclic form of a single monosaccharide (pyranose or furanose) or a single linkage. Due to the very small differences in isomeric confirmation between these four monosaccharides, a modified gradient was applied (35 to 40% B over 170 mins) (**Figure 5**). The four standards were analyzed separately (**Figure 5A-D**) or as a mixture (**Figure 5E**) and demonstrated nearly baseline separation using the modified gradient program. Ofthe four isomeric O-glycan structures, the pyranose structures eluted over 20 mins earlier than furanose-containing isoforms. As described previously, the observed retention times were converted to reverse phase glucose units (RP g.u.) using the linear regression fitting from dextran ladder runs before and after the sample queue using the same gradient program (**Figure 5F**).

**Figure 4.**
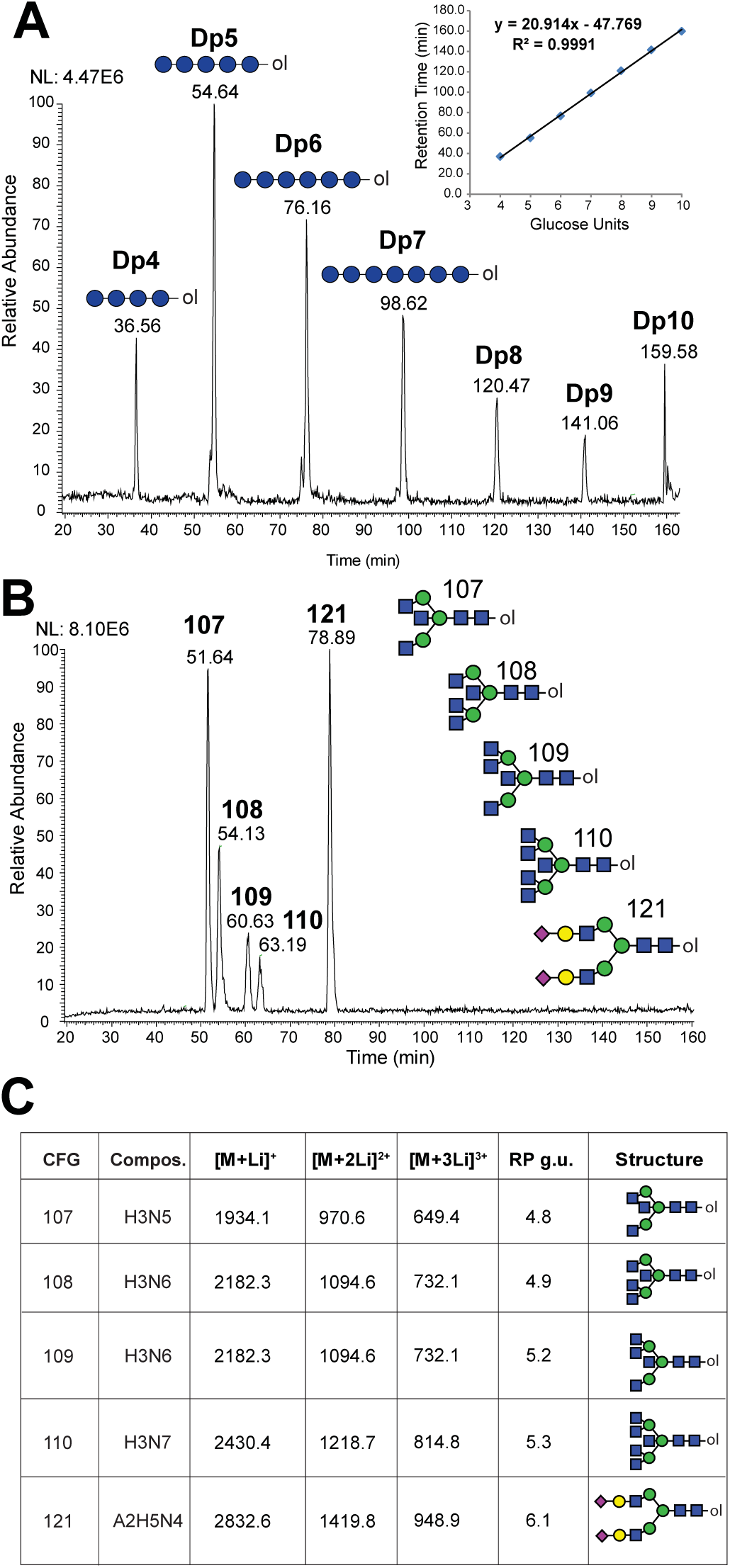
Elution of permethylated glycans expressed in terms of Glucose Units. (**A**) Reduced and ^13^C-permethylated Dextrans (isomaltooligosaccharide series, DP4 to 10) were separated using a PepMap Acclaim C18 column in 1 mM lithium acetate-containing mobile phases (45-70% B over 150 mins). Extracted ion chromatogram for each predicted mass is shown (within 10 ppm). Retention times of DP standards, run both before and after sample acquisition queue, were averaged and plotted using a linear regression fitting. Error bars represent standard deviation (See *inset).* (**B**) Reduced and ^13^C-permethylated CFG N-Glycan standards (107-110, 121) were analyzed as described in Panel A. (**C**) Summary table of analyzed CFG standards with retention times converted to reversed phase glucose units (RP g.u.) using linear regression fitting shown in Panel A, *inset*).

**Figure 5.**
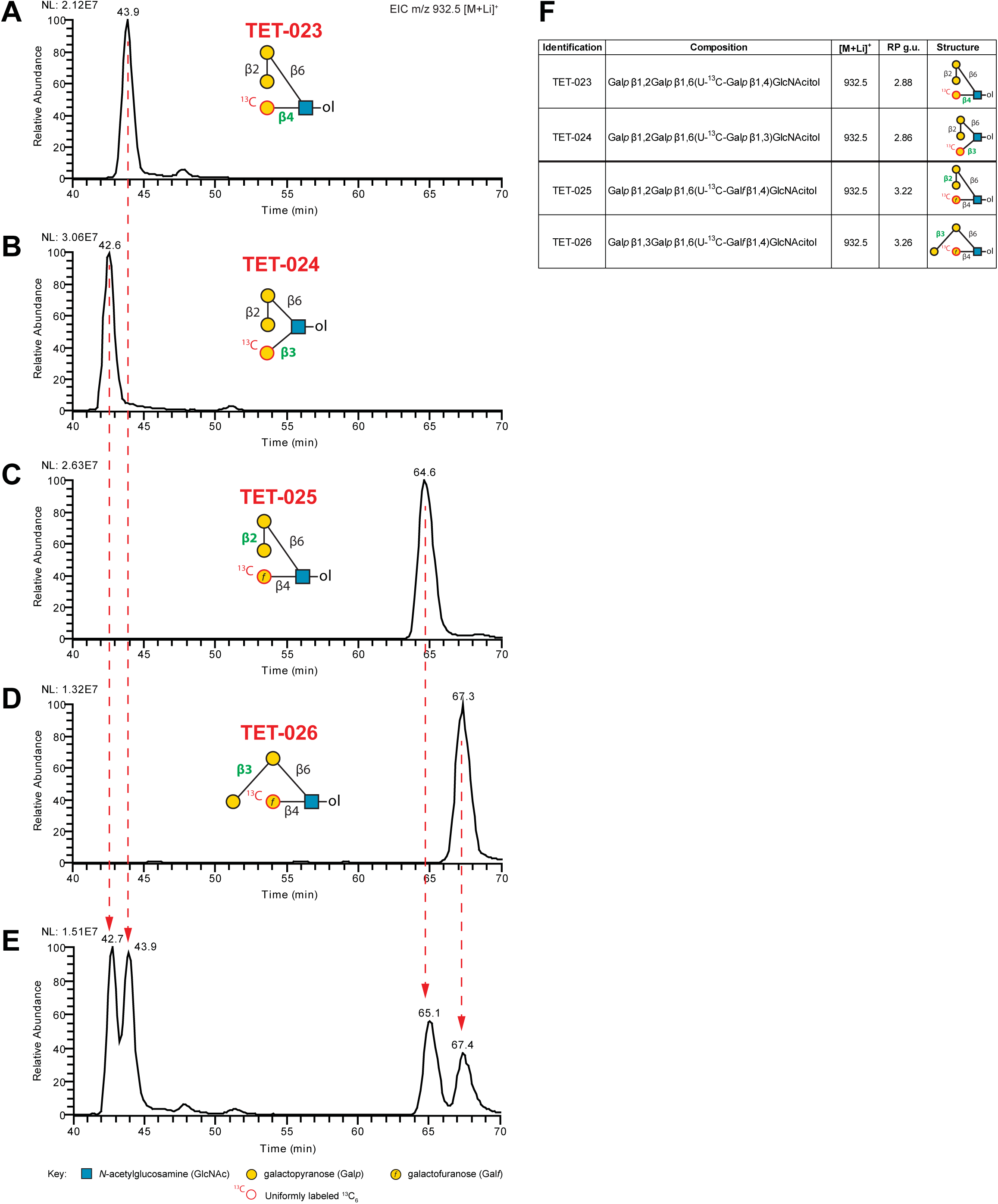
LC-NSI-MS of Permethylated Isomers. (**A**) Extracted ion chromatograms showing individual (**A-D**) or a mixture (**E**) of four O-glycan tetrasaccharide isomers that only differ by the cyclic form of a single sugar [pyranose (p) or furanose (f)] or a single linkage (differences indicated in green). Separation was performed using a PepMap Acclaim C18 column in 1 mM lithium acetate-containing mobile phases with a modified gradient (35-40% B over 170 mins) and detected using a Thermo Velos Pro ITMS. (**F**) Summary table of isomers used with retention times converted to reversed phase glucose units (RP g.u.; referenced to ^13^C-permethylated isomaltooligosaccharide standards separated using the same gradient).

### Characterization of N-glycans released from recombinant HIV gp120 expressed in glycoengineered Pichia pastoris

The data described above demonstrates the ability of our method to resolve N- and O-glycan structures derived from abundant glycoproteins or prepared as synthetic standards. To assess the applicability of our method for detecting biologically relevant glycosylation patterns, we analyzed N-glycans released from recombinantly expressed HIV gp120 envelope glycoproteins harvested from six different glyco-engineered strains of *Pichia pastoris* (13, 14). Each of these strains lack glycosyltransferase activities that result in the generation of restricted cellular N-glycan profiles predicted to be to predominated by specific high-mannose or complex type glycans. After release, reduction, and permethylation, N-glycans from each gp120 preparation were first separated and analyzed individually, demonstrating that the detected N-glycans are consistent with the targeted glycosyltransferases in each glyco-engineered strain (**Figure 6, A-F**). For example, N-glycans of the composition Hex_3_HexNAc_2_-ol were only detected in the strain engineered to generate Man_5_GIcNA_2_ structures on gp120 glycoprotein (**Figure 6, A**) and Hex_5_HexNAc_2_-ol in the Man_5_GlcNAc_2_ engineered strain (**Figure 6, B**). A similar pattern was observed in engineered strains producing only Man_8_GlcNAc_2_ and Man_9_GlcNAc_2_ as well as Hex_8_HexNAc_2_-Hex_10_HexNAc_2_ structures (**Figure 6, C-D**). *Pichia* strains engineered to synthesize N-glycan structures with terminal galactose residues or terminal sialic acid modifications also exhibited Man_5_GlcNAc2 structures in addition to the expected complex type structures (**Figure 6, E-F**). To assess the capacity of our LC-MS system to resolve the range of glycans released from these expression platforms, we mixed equal amounts of each sample and analyzed as described above. We were able to accurately detect and resolve all of the various N-glycan structures derived from the recombinant gp120 glycoforms that were detected when each were analyzed individually (**Figure 6, G**). Furthermore, we applied an automated MS^n^ fragmentation workflow which targeted the neutral loss of permethylated sialic acid (Δm/z = 375) to trigger product dependent MS^3-5^ acquisitions. Since lithium was included in the LC separation mobile phases, this workflow revealed that the terminal sialic acids of the N-glycan structures released from these gp120 preparations are a2,6-linked (**Figure 6, H and I**).

**Figure 6.**
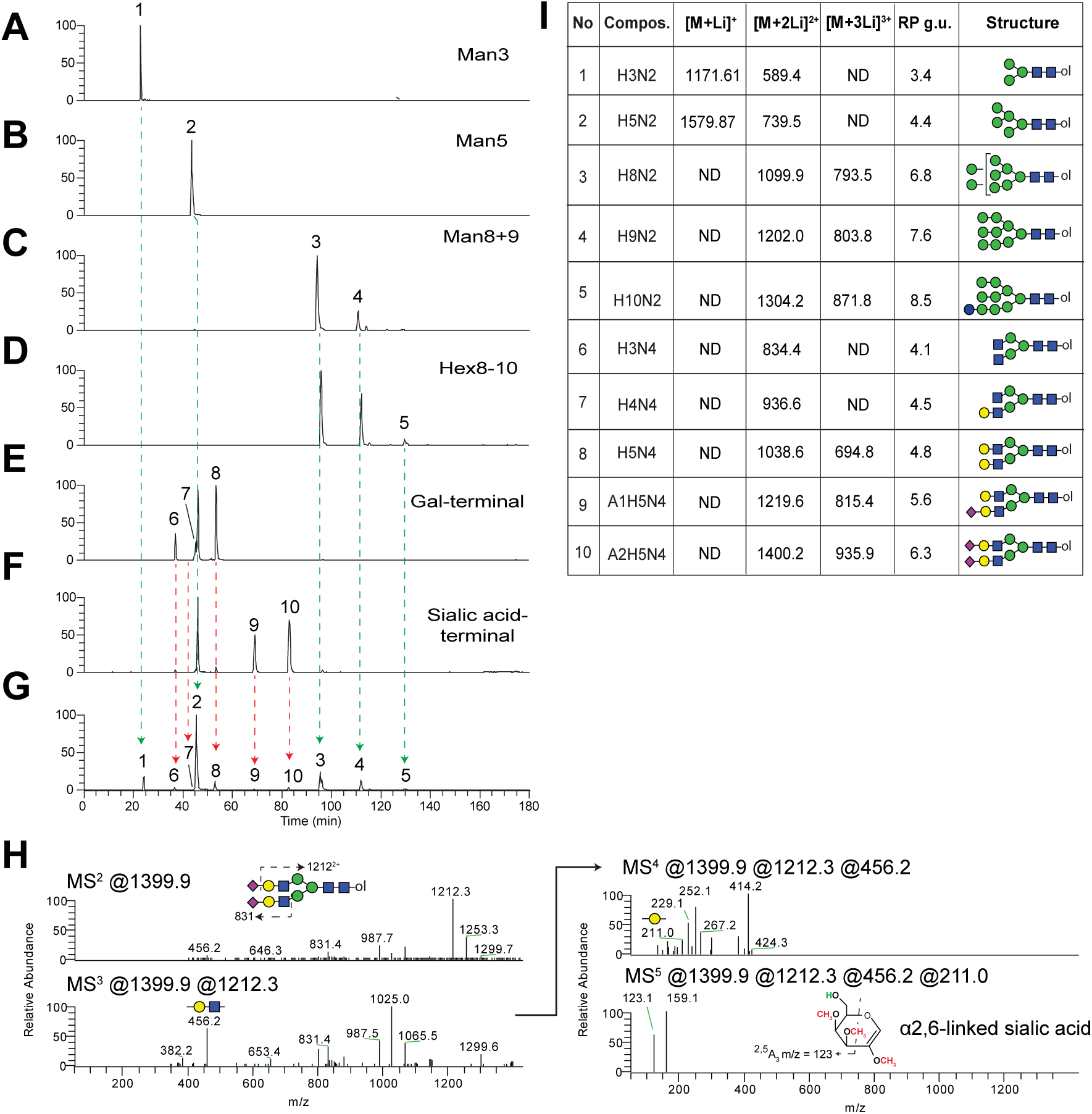
LC-NSI-MS^n^ analysis of N-glycans released from recombinant gp120 expressed in *Pichia pastoris* strains defective in N-glycosylation machinery. (**A**) Extracted ion chromatograms of all predicted PNGaseF-released, reduced, and permethylated N-glycans from various gp120 expressing cell lines, either individually (**A-F**) or as a mixture (**G**). (H) Automated, intelligent MS^n^ fragmentation defines sialic acid linkage position in realtime from Structure #10. (**I**) Summary table of gp120 N-glycans with retention times converted to reversed phase glucose units (RP g.u.; referenced to ^13^C-permethylated isomaltooligosaccharide standards separated using the same gradient).

### Analysis of O-glycan isomers from biological samples

O-linked glycans linked to Ser/Thr residues can be divided into various subtypes depending on the initiating monosaccharide linked directly to the polypeptide backbone. The most abundant class is the mucin-type O-GalNAc-initiated structures. Other types of O-glycans include O-Mannose, O-Fucose, O-Glucose, O-GIcNAc, O-Xylose, all of which have diverse biological functions (23, 24). Previous studies have documented the detection of both O-GalNAc and O-Mannose-initiated disaccharides, Gaiβ3GalNAc and GlcNAcβ2Man, respectively, in mouse brain (12, 25). In direct infusion MS, these glycans are detected at the same m/z since they have the same Hex_1_HexNAc_1_ composition. While MS/MS fragmentation of the permethylated disaccharides can detect the presence of these two glycans, the fragmentation patterns of the isolated ions can be complicated as they are a mix of the two species and quantification of each individual disaccharide is problematic. To identify the presence or absence of Hex_1_HexNAc_1_ isomers in mouse brain, we analyzed the released and permethylated glycans by our LC-MS workflow (**Figure 7, A**). An EIC ofthe expected singly-charged, lithiated structures (*m/z* 518.3) revealed two peaks of differing relative abundance at 17.8 and 19.2 min. MS2 fragmentation of m/z 518 at the respective times is consistent with the HexNAc-initiated disaccharide (O-GalNAc) eluting earlier than the Hexose-initiated structure (O-mannose), as determined by unique fragments at the reducing terminus (**Figure 7, A, insets**).

**Figure 7.**
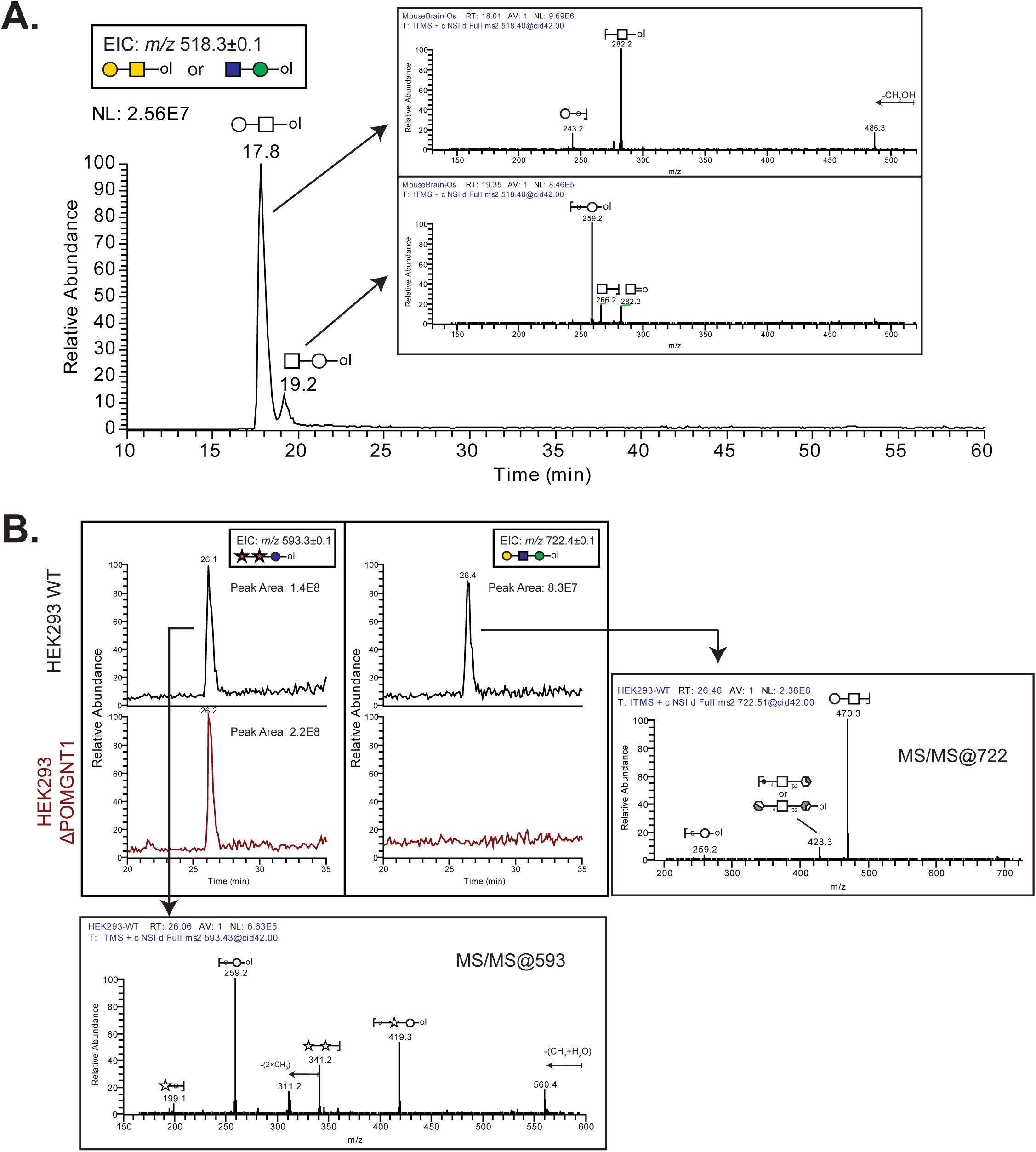
Analysis of reduced and permethylated O-glycans released from mouse brain proteins and human cell lines defective in the O-mannosylation pathway. (**A**) Extracted ion chromatogram of HexiHexNAci structures released from mouse brain proteins indicates two structural isomers. MS2 fragmentation of m/z 518.3 at the indicated retention times are shown as insets. (**B**) LC-MS analysis of O-glycans released from HEK293 WT and APOMGNT1 mutant cell lines. Respective EICs and annotated MS2 spectra are shown.

We also analyzed O-glycans harvested from cells that were glyco-engineered to be deficient in the O-Mannosylation pathway implicated in congenital muscular dystrophies (26-28). The glycosyltransferase POMGNT1 (Protein O-Linked Mannose N-Acetyl-glucosaminyltransferase 1) transfers β-linked GIcNAc to an underlying α-mannose linked to SerorThrto generate Core Mi-type glycans which can be further extended by B4GALT1 with β4-Iinked galactose (29). O-glycans were released from glycoproteins harvested from HEK293 wildtype cells or HEK293 cells harboring a Crispr-Cas9-mediated global disruption of the P0MGNT1 locus (30). As a control, a non-O-Mannose structure (Xylα3Xylα3Glc-ol), which modifies a consensus found in the EGF repeats of a relatively small number of proteins (31-33), was detected (m/z 593.3 [M+Li]^+^) in both samples (**Figure 7B**). An EIC of m/z 722.1 ([M+Li]^+^) which corresponds to the composition Hex_2_HexNAc_1_ (O-mannose Core M1 structure, Gaiβ4GlcNAcβ2Man-ol) demonstrates the presence ofthe glycan in HEK293 WT but not in the POMGNTI-deficient cell line.

## DISCUSSION

The field of glycomics has historically been eager to engage new analytic platforms based on their promise to deliver greater sensitivity and enhanced detection of structural features (34-37). Despite tremendous advances in various analytic technologies, it remains true that no single technique, besides NMR, is capable of routinely and completely defining glycan structures. Thus, the field will continue to benefit from the development of an expanding array of analytic approaches. We have combined two powerful technologies, glycan permethylation and liquid chromatography, in a manner that should be adoptable by analytic laboratories familiar with standard LC-MS/MS proteomic techniques. To facilitate adoption of our nano-LC-NSI-MS^n^ approach, we have optimized run conditions and buffer compositions for analysis of N-linked or O-linked glycans. Using these conditions, we achieved baseline separations of challenging isomeric glycans and also demonstrated the power of on-the-fly MS^n^ to distinguish structural features such as the linkage position of sialic acids.

Other investigators have previously described LC-based separations of glycans and, more particularly, of permethylated glycans (3, 9, 10). These approaches are all valuable contributions to the arsenal of glycomic technology. In many cases, these techniques make use of glycan standards to define the retention times of known structures, generating a database of RT values that, when combined with accurate mass detection, can provide confident identification of components in complex mixtures. Glycans with novel RT values can be at least partially characterized using these methods by applying off-line exoglycosidase digestions (sequencing) and re-analysis (37). Our approach also takes advantage of RT and accurate mass to identify glycans, but additionally includes the opportunity to capture MS^n^ data on-the-fly to help assign structures with novel RT values in a single analytic run. In addition, we have enhanced this capability by including lithium in the mobile phase which facilitates the generation of cross-ring fragments indicative ofspecific structural features (5).

We optimized the lithium concentration in the mobile phase so that we detect glycans as a single adduct species. This simplification allows quantification to be done from peak areas detected in MS1 for a single adduct, enhancing sensitivity and resolution of glycan species across the chromatogram. Our method incorporates the inclusion of RT standards (DP4-10) which allows the calculation of relative retention times in Glc units (RP-g.u.), facilitating cross-platform and run- to-run identification of glycans of interest. Additionally, these standards provide a means to quantify glycan abundance against a known amount of glycan. We have previously demonstrated that permethylated glycans yield nearly the same signal intensity regardless of their mass or complexity, unless modified with sulfate or phosphate (4). Thus, known amounts of DP standards can serve not only as RP-g.u. references but also as quantification standards to provide calculation ofabsolute abundance.

Previously published glycomic studies from our laboratories and many others, more than can be reasonably cited, have relied heavily on direct infusion of permethylated glycans. These experiments, while valuable for in-depth structural analysis, require the presence of the operator for sample load, run, and subsequent wash before load of next sample. The nanoLC-NSI-MS^n^ method we describe here, like other LC-based methods, is amenable to autosampling, greatly increasing analytic throughput. Furthermore, data collection is almost entirely automated compared to more labor-intensive direct infusion experiments. This marked benefit also highlights a bottleneck in all glycomic analysis, but especially for LC-based analysis, namely, facile analysis of large datasets. Software platforms capable of handling and annotating large glycan datasets generated by nanoLC-NSI-MS^n^ approaches are only in their infancy (38). We made use of our in-house developed GRITS Toolbox to sieve through the data by providing structural candidates based on MS/MS data (18). However, neither this platform nor any other is currently capable of integrating MS^n^ data for permethylated glycans across an entire LC run. Furthermore, software packages developed for LC-based proteomic applications are not ideal for interpreting LC of permethylated glycans because the underlying assumptions regarding isotopic distributions of amino acids are not transferable to permethylated monosaccharides.

Fully automated annotation packages for handling permethylated glycan LC-separation data is a significant need for expanding the adoption of this powerful approach for glycomics. Here, we have provided a useful and robust separation platform that will facilitate the generation of data essential for developing and testing such tools. In the meantime, as these new software tools are developed and implemented, the expanded ability of our nanoLC-NSI-MS^n^ platform to resolve and quantify glycan isomers, combined with the capacity for high-throughput analysis, will provide new opportunities for investigating the diversity of glycosylation at scales capable of answering biologically important questions.

## ABBREVIATIONS

nanoLC-NSI-MS: nanoflow Liquid Chromatography-Nanospray Ionization-Mass Spectrometry
MS^n^: multi-dimensional ion fragmentation
ACN: Acetonitrile
MeOH: Methanol
AcOH: Acetic Acid
LiOAc: Lithium Acetate
DP: Degree of Polymerization
ITMS: Ion Trap Mass Spectrometer
CID: Collision Induced Dissociation
NaBH_4_: Sodium Borohydride
NaOH: Sodium Hydroxide
EIC: Extracted Ion Chromatogram
RP: Reverse Phase
SPE: Solid phase extraction
Mel: methyl iodide

## FOOTNOTES

### Author Contributions

S.K., M.O.S, M.T. and L.W. designed research; S.K. and M.O.S performed research; S.L. contributed key reagents; S.K., M.O.S, M.T. and L.W. analyzed data; S.K., M.O.S, M.T. and L.W. wrote the paper.

## Acknowledgements

The authors wish to gratefully acknowledge Brent Weatherly for his graciously given advice, encouragement, and assistance regarding computational analysis and informatic considerations over the course of this project and others, before his untimely death. We also thank Rob Bridger for his technical support of our mass spectrometry instrumentation, Dr. Christopher M. West (University of Georgia) for providing the *T. cruzi* O-Glycan isomeric standards, and Dr. Henrik Clausen (University of Copenhagen) for providing HEK293 cell lines. We are grateful for technical advice from Drs. Christa Feasley, Julian Saba, and Sergei Snovida (ThermoFisher Scientific). Four N-linked glycan standards (standards 107 to 110) were received from the Consortium for Functional Glycomics (CFG) and an additional N-linked glycan standard (standard 121) was received as a gift from Dr. James Paulson (The Scripps Research Institute). The instrumentation utilized in this analysis was provided by ThermoFisher in partial support of the ThermoFisher Center of Excellence for Glycomics and Glycoproteomics at the CCRC, Lance Wells and Michael Tiemeyer, Co-Directors.

## FIGURE LEGENDS

**Supplement 1. Reduction of N-glycans prior to permethylation eliminates peak-splitting during LC-separation.** N-glycans released from RNAse B by PNGase F digestion were permethylated and subjected to PepMap Acclaim C18 LC separation in 1 mM lithium acetate-containing mobile phases and detected using Thermo Velos Pro ITMS. Extracted Ion chromatograms of predicted RNAse B permethylated Man5-Man9 structures demonstrate that non-reduced N-glycans tend to peak-split due to the presence of α- and β-anomeric configurations prior to permethylation (**A**), while glycans that are reduced prior to permethylation do not (**B**).

**Supplement 2. Reproducibility of standard performance. (A**) Reduced and ^13^C-permethylated Dextran (isomaltooligosaccharide series, DP4 to 9) was separated using a PepMap Acclaim C18 column in 1 mM lithium acetate-containing mobile phases (45-70% B over 150 mins). Extracted ion chromatogram for each predicted mass is shown (within 10 ppm). (**B**) Rerun of Dextran after 25 days of continuously running samples.

**Supplement 3. Limit of Detection.** Reduced and ^13^C-permethylated isomalto analytical standards DP4-6 were analyzed by LC-NSI-MS at different amounts ranging from 15.625 to 1000 femtomoles injected and EICs were evaluated based on (**A**) Peak apex retention time, (**B**) Peak Area, and (**C**) Peak Height.

